# Predicted mechanistic impacts of human protein missense variants

**DOI:** 10.1101/2024.05.29.596373

**Authors:** Jürgen Jänes, Marc Müller, Senthil Selvaraj, Diogo Manoel, James Stephenson, Catarina Gonçalves, Aleix Lafita, Benjamin Polacco, Kirsten Obernier, Kaur Alasoo, Manuel C. Lemos, Nevan Krogan, Maria Martin, Luis R. Saraiva, David Burke, Pedro Beltrao

## Abstract

Genome sequencing efforts have led to the discovery of tens of millions of protein missense variants found in the human population with the majority of these having no annotated role and some likely contributing to trait variation and disease. Sequence-based artificial intelligence approaches have become highly accurate at predicting variants that are detrimental to the function of proteins but they do not inform on mechanisms of disruption. Here we combined sequence and structure-based methods to perform proteome-wide prediction of deleterious variants with information on their impact on protein stability, protein-protein interactions and small-molecule binding pockets. AlphaFold2 structures were used to predict approximately 100,000 small-molecule binding pockets and stability changes for over 200 million variants. To inform on protein-protein interfaces we used AlphaFold2 to predict structures for nearly 500,000 protein complexes. We illustrate the value of mechanism-aware variant effect predictions to study the relation between protein stability and abundance and the structural properties of interfaces underlying *trans* protein quantitative trait loci (pQTLs). We characterised the distribution of mechanistic impacts of protein variants found in patients and experimentally studied example disease linked variants in FGFR1.

## Introduction

Over 16 million protein missense variants have been found in the 730 thousand human exomes currently integrated into the gnomAD (gnomad.broadinstitute.org) resource (Chen *et al*, 2024). Among the 454,787 individuals with exomes sequenced in the UK Biobank, the typical distribution of variants per individual is on the order of 9,292 missense variants per individual, with the vast majority being rare in the population and around 23% predicted to be deleterious (Backman *et al*, 2021). It is increasingly clear that such rare variants can contribute significantly to human trait variation and disease but only a very small fraction of these have any functional annotations. For example, around 49 thousand missense variants have a functional annotation based on public repositories (Landrum *et al*, 2014; Whirl-Carrillo *et al*, 2021; Wright *et al*, 2023; Chang *et al*, 2018).

Experimental characterization of protein missense variants is now possible to scale up, given the development of deep mutational scanning (DMS) experiments (Fowler & Fields, 2014). These methods allow for the measurement of impact of all possible missense variants on protein function as measured in pooled screens. These DMS screens are primarily designed to identify variants that impact on the protein function as a whole but specific assays have also been designed to select for variants relevant for protein stability, binding or catalytic activity (Faure *et al*, 2022; Tsuboyama *et al*, 2023; Geck *et al*, 2022). In parallel to experimental approaches there have been important recent developments in artificial intelligence methods to predict the impact of missense variants on protein function. There has been a long history of development of such approaches, including methods that rely on evolutionary information as well as protein structure-based methods. While methods like SIFT (Ng & Henikoff, 2003) and Polyphen (Adzhubei *et al*, 2010) continue to be very actively used, more recent neural network-based methods such as EVE (Frazer *et al*, 2021), ESM1b (Brandes *et al*, 2023) and AlphaMissense (AM) (Cheng *et al*, 2023) represent very significant improvements that have been shown to sometimes compete with experimental measurements from DMS experiments in accuracy when being used to discriminate between pathogenic and benign variants (Livesey & Marsh, 2023).

The developments in experimental and computational methods have been primarily driven by a need to identify protein missense variants that are detrimental to protein function. While there may still be more room for improvements, one pressing challenge that remains is to develop methods that can assign a biochemical mechanism for the impact of the protein variants. There are multiple ways by which a missense variant could result in a deleterious effect, including a change in protein stability, enzymatic activity, binding to other proteins or other biomolecules such as DNA/RNA, small molecules or lipids. Knowledge of such effects may be relevant for diagnostic purposes since proteins have multiple functions and the disruption of different functions may result in different organismal traits. More importantly, knowledge of the disrupted mechanism may be critically relevant for the development of therapies.

Protein structure information is very useful in the study of the mechanist impact of missense variants. Structures can be used to infer whether a missense variant can result in changes in protein stability and interactions using a growing array of computational approaches (Delgado *et al*, 2019; Blaabjerg *et al*, 2023; Gerasimavicius *et al*, 2023). The development of AlphaFold2 (AF2) and related methods indicates that we can use structural data at proteome-wide scale to infer mechanisms of disruption by missense variants. Here, we have used AF2 predicted structures to infer the impact on protein stability for nearly all human positions with a high confidence AF2 prediction, covering over 200 million variants. We have identified small-molecule binding pockets in the human proteome and used AF2 to predict the structures of nearly 500,000 protein-protein interactions. The predicted residues in pockets or at protein-protein interfaces are enriched in disease linked variants indicating these are likely to be important functional residues. We use this resource to study the relation between protein stability and abundance for cancer mutation. Using the interface models we studied the distribution of predicted interface regisions and how the structural properties of protein-protein interfaces relate to the coordinated regulation of protein levels. We provide a proteome-wide set of functional annotations for all possible missense variants accessible through a web resource.

## Results

### Proteome-wide prediction of protein stability changes

The availability of protein structures predicted for the full human proteome allows for the large-scale structure-based inference of protein stability changes. We have previously shown that the FoldX algorithm can be applied to predict stability changes for protein variants based on AlphaFold2 (AF2) in regions of median to high confidence (Akdel *et al*, 2022). Recent benchmarks indicate that FoldX remains one of the top performing stability predictors (Gerasimavicius *et al*, 2023). Here, we used FoldX to predict stability changes across all human AF2 predicted structures with confidence scores pLDDT>70, corresponding to 208 million predicted scores. We considered that missense variants that are likely to be destabilising would serve as adequate independent benchmarks for testing of state-of-the art pathogenicity predictions. Based on our predictions, we identified 8019 human proteins with over 1000 variants predicted to be neutral or destabilising based on structural information and used these to compare two recently developed variant effect predictors (VEPs) (ESM1b and AlphaMissense) (**Fig 1A**). Based on these benchmarks we observed that AlphaMissense outperforms ESM1b with only a small fraction of genes showing better predictions using ESM1b.

**Fig 1.**
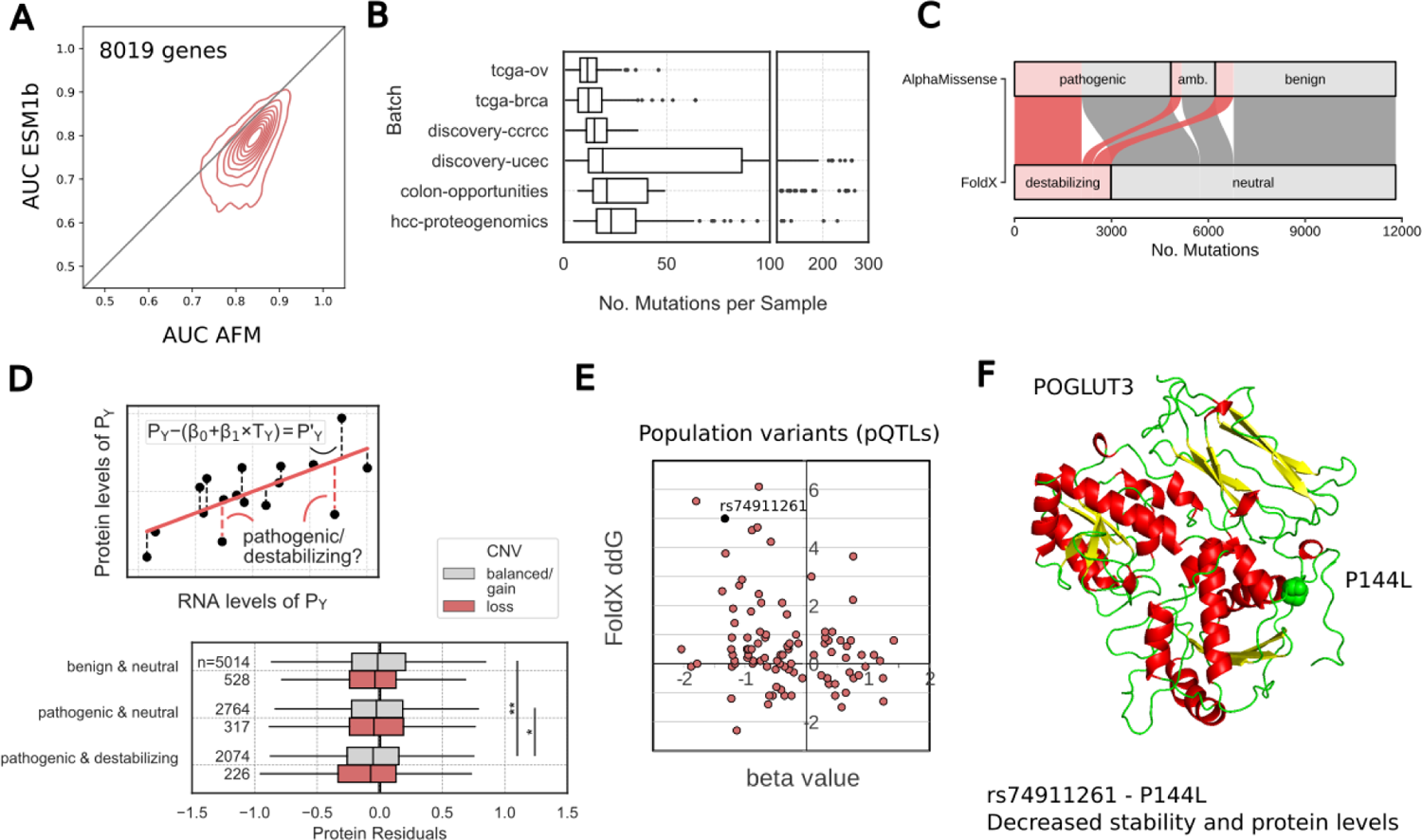
Genome-wide prediction of protein stability changes. **A.** AlphaMissene and ESM1b scoring of variants predicted to destabilise or not protein structures for 8019 human proteins with over 1000 putative destabilising and 1000 neutral variants. For each protein we used both predictors to attempt to discriminate between putative destabilising and neutral variants summarising each protein result with the value for the area under the ROC curve (AUC). The distribution of AUCs for both predictors in the 8019 proteins is shown as a density plot. **B.** Mutational burden of cancer samples by tissue. The overall median number of missense mutations per sample was 17. **C.** Sankey plot of combinations of missense variant effects as predicted by FoldX (stability) and AlphaMissense (pathogenicity). 11802 mutations could be annotated using both AlphaMissense and FoldX. 69% of predicted destabilising variants were also predicted pathogenic, but only 43% of predicted pathogenic mutations were also predicted destabilising. **D.** upper plot: illustration of the method for estimating residual variance of protein abundance after accounting for the expected level of protein abundance expected by their mRNA level (see Methods). We ask if negative residuals are associated with destabilising and/or pathogenic variants. Lower plot: Analysis of protein residuals in genes carrying missense mutations for different predictions of pathogenicity (AlphaMissense) and stability (FoldX). Grey box plots represent cancer samples in which at least one other, non-mutated allele of a gene is present which could potentially compensate for the deleterious effect of the mutated allele on the affected protein. Red box plots represent cancer samples with copy number loss in the affected gene, where such compensation would not be possible. **E.** Scatter plot of: x-axis - beta values for significant associations of missense variants with changes in the corresponding protein levels (cis pQTLs), y-axis - changes in stability predicted for the same missense variants using FoldX (ΔΔG in kcal/mol). **F.** Predicted structure for Protein O-Xylosyltransferase (POGLUT3, KDELC2) illustrating the positions for the P144L variant (rs74911261).

Not all pathogenic variants are expected to lead to destabilisation and it is thought that pathogenic variants that destabilise structures are more likely to result in protein degradation (Gerasimavicius *et al*, 2023). To test this on a large scale we compiled a multi-omics cancer dataset where data is available for the genome, transcriptome and proteome in 644 samples from 6 tumour types, carrying typically 17 missense mutations per sample (**Fig 1B**). We used AlphaMissense to classify 11802 cancer missense variants found in these samples into those that may disrupt protein function and we then used FoldX scores to classify them into those that are predicted to be destabilising (**Fig 1C**). From the variants that are predicted to result in changes in stability 69% were considered to be deleterious by AlphaMissense (**Fig 1C**). However, only around 43% of AlphaMissense variants predicted to be deleterious were likely to impact on protein stability, indicating the importance of considering other functional roles of protein residues.

We then used the mRNA and protein abundance samples to estimate the protein levels in a sample that deviates from the expected correlation between changes in mRNA and protein levels (**Fig 1D**). This deviation is expected to include sample specific changes in translation, degradation, technical noise or other factors. We hypothesise that samples carrying variants predicted to destabilise a protein may show lower abundance for the destabilised proteins. We tested if different types of missense mutations were more likely to correspond to a loss in protein levels. On average, we observed that variants that were predicted as pathogenic - based on AlphaMissense - were more likely to be associated with a decrease in protein levels only when they were also predicted to be destabilising to the protein structure (**Fig 1D**). This was particularly true for mutated proteins that had only 1 copy of the gene present (**Fig 1D**).

Several human population variants have been linked to changes in protein levels in the plasma based on quantitative trait locus (QTL) analysis (Tambets *et al*, 2024; Sun *et al*, 2018). We used our stability prediction to annotate a set of 98 protein missense variants that have been statistically associated with changes in the corresponding protein levels, so called cis pQTLs. Compared to all variants, those that are predicted to lower the stability of the protein (ΔΔG>1.5 kcal/mol) were enriched in loss of protein levels (negative beta values, p-value<0.05 fisher’s exact test, **Fig 1E**). As an example, the variant rs74911261 is a human population variant that results in a missense change between proline and leucine at position 144 of the Protein O-Xylosyltransferase (POGLUT3, also known as KDELC2), involved in protein O-linked glycosylation. The proline to leucine mutation is predicted to be destabilising and is statistically linked with a lower amount of this protein measured in plasma. The same SNP is also linked to increased risk of breast, renal and prostate cancer suggesting that POGLUT3 protein stability is a risk factor for cancer. In total we identified 13 population variants that are predicted to be destabilising (ddG>2) that are also associated with reduced protein levels.

### FGFR1 patient variants linked with congenital hypogonadotropic hypogonadism

We next studied protein variants on a cohort of patients suffering from congenital hypogonadotropic hypogonadism (CHH), a rare disorder caused by a deficiency in Gonadotropin releasing hormone (GnRH). In this context, FGFR1 signalling plays a critical role in the development and function of the hypothalamic-pituitary-gonadal (HPG) axis, which is essential for normal reproductive function. Mutations in the FGFR1 gene can lead to impaired signalling, which is associated with CHH (Gonçalves *et al*, 2015). FGFR1 signalling is involved in the embryonic development of GnRH neurons and their migration to the hypothalamus. Disruptions in this pathway can result in the failure of GnRH neurons to reach their proper location or function correctly, leading to the clinical manifestations of CHH. Understanding the role of FGFR1 signalling in CHH is crucial for developing targeted therapies to address the underlying causes of this disorder (Falardeau *et al*, 2008).

We identified in this cohort seven protein variants in FGFR1, including one indel, one synonymous and five missense variants. From these variants, we selected three examples for functional studies, a missense variant predicted to be both pathogenic and destabilising (M719V), a missense variant predicted to be pathogenic but not destabilising (S96C), and the frameshift (Y654fs). We evaluated the effects of three FGFR1 variants on total protein levels, membrane localization, cell growth and the activation of downstream signalling pathways activation of several downstream signalling pathways, including the MEK-ERK, AKT, and mTOR pathways in GT1-7 mouse hypothalamic neuronal cells, with wild-type (WT) FGFR1 serving as a reference (**Fig 2A-E**). In line with the stability predictions, the M750V variant, unlike the S129C variant, exhibited reduced total protein levels, a trend also noted in the frameshift mutant (**Fig 2A**). Notably, all three mutants displayed decreased membrane localization (**Fig 2B**). Additionally, cell growth was decreased in cells transfected with mutant FGFR1 compared to those transfected with WT FGFR1 (**Fig 2C**). Subsequent experiments showed that, upon FGF2 treatment to stimulate the FGFR1 receptor, cells expressing mutant FGFR1 variants were unable to phosphorylate critical proteins in the MEK, ERK, AKT, and mTOR pathways, unlike cells expressing WT FGFR1 (**Fig 2D,E**). This impairment in pathway activation may be linked to the reduced membrane presence of the mutant FGFR1 proteins, highlighting a possible explanation for the observed functional impairments. Impaired activation of the MEK-ERK, AKT, and mTOR signalling pathways due to mutant FGFR1 membrane expression can lead to compromised cellular functions, contributing to the pathogenesis of CHH. This may manifest as decreased proliferation and survival of GnRH neurons, ultimately resulting in reduced GnRH production and the associated reproductive and hormonal deficiencies characteristic of CHH (Eswarakumar *et al*, 2005; Turner & Grose, 2010; Pitteloud *et al*, 2006).

**Fig 2.**
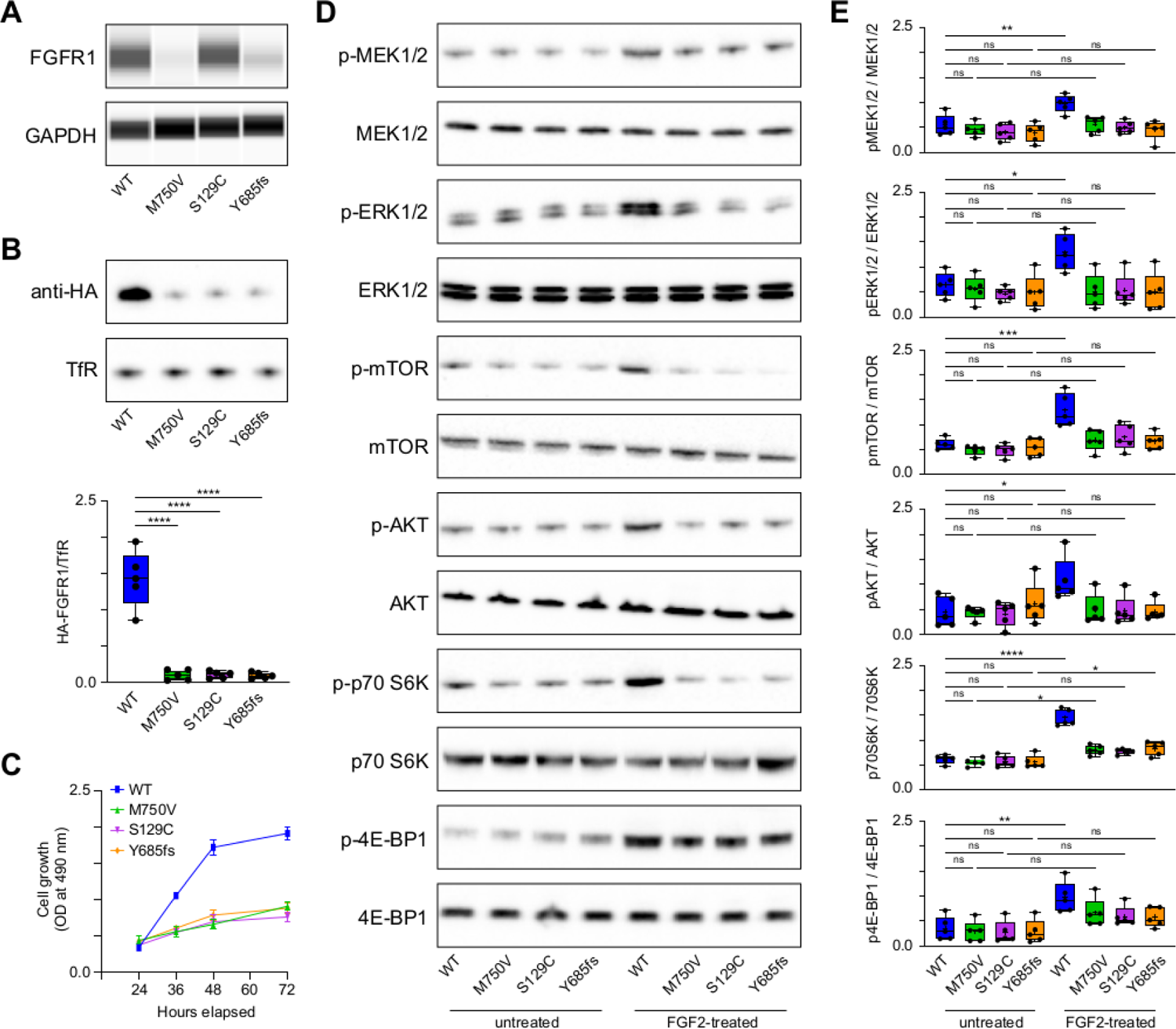
FGFR1 mutations associated with congenital hypogonadotropic hypogonadism. **A.** FGFR1 Expression in GT1-7 Cells: Analysis of HA-tagged wild-type (WT) and mutant FGFR1 expression in GT1-7 cells transiently transfected with the respective constructs. Cells were lysed 48 hours post-transfection, and the lysates were analysed using a ProteinSimple WES automated western blot system with a 12-230 kDa separation module to assess the expression levels of WT and mutant FGFR1. **B.** Cell Surface Expression of FGFR1: Evaluation of cell surface expression of WT and mutant FGFR1 in GT1-7 cells transfected with HA-tagged constructs. After cell surface biotinylation, proteins were isolated, resolved by SDS-PAGE, and detected using an anti-HA antibody on western blots. Transferrin receptor (TfR) serves as a loading control to ensure equal protein loading and to validate the efficiency of the biotinylation process. **C.** Cell Growth Assay: Assessment of cell growth in GT1-7 cells transiently transfected with WT or mutant FGFR1 using the MTT assay. Cell growth was measured at 24, 36, 48, and 72 hours post-transfection to evaluate the impact of FGFR1 variants on cell proliferation. **D.** Activation of signalling Pathways: Analysis of the activation of MEK1/2, ERK1/2, mTOR, AKT, p70S6 kinase, and 4E-binding protein 1 (4EBP1) in GT1-7 cells transfected with WT or mutant FGFR1. Cells were stimulated with 100 ng/ml FGF for 60 minutes, 36 hours post-transfection. Western blotting was performed to detect total and phosphorylated (phospho) forms of MEK1/2, ERK1/2, mTOR (p-mTOR), AKT (p-AKT), p70S6 kinase (p-p70S6K), and 4EBP1 (p-4EBP1). **E.** Quantification of Phosphorylated Proteins: The phosphorylated MEK1/2, ERK1/2, mTOR, AKT, p70S6 kinase, and 4EBP1 blots were quantified using their respective total proteins to assess the activation of these signalling pathways in response to FGF stimulation in cells expressing different FGFR1 variants. This quantification provides insights into the impact of FGFR1 variants on the activation of key signalling pathways involved in cell growth and proliferation.

### Binding site predictions facilitate interpretation of (pharmaco)-genomic variants

Only around one third of likely pathogenic cancer variants were also predicted to result in changes in protein stability, indicating the importance of annotating the functional mechanisms of other protein residues (**Fig 1C**). Small-molecule binding sites form another set of important functionally relevant residues, along with residues that contribute to the stability of the structure and protein-protein interfaces. Here we again made use of the AlphaFold2 predicted structures to perform a proteome-wide search for putative small-molecule binding pockets. We have seen before that AlphaFold2 predictions can result in spurious pockets forming in regions of high uncertainty (Akdel *et al*, 2022). We defined a pocket scoring scheme that includes the residue level uncertainty estimates from AlphaFold2 (see Methods, **Fig S1**). We ranked the 547,401 predicted pockets according to this score and can show that known small-molecule binding sites consistently rank at the top (**Fig 3A**, AUC=0.87). Given that the known binding sites are derived from experimental structures we also tested the method’s capacity to identify pockets in enzymes that have no experimental structure or high identity to solved structures. The lack of any homology to existing structures indicates that these proteins and their pockets could not have been in the training data for AlphaFold2. We find that pockets in these proteins were also consistently ranked at the top of our pocket score (**Fig 3A**, AUC=0.82). Based on these observations we selected the top 109,599 as high confidence pockets found across nearly all human proteins. Most proteins tend to have a small number of predicted pockets (**Fig 3B**), with 2,420 of 20,296 (11.92%) having one predicted pocket and around half of the proteome (9,813 of 20,296; 48.35%) having 3 or less. Only 26.6% of these pockets could have been predicted before AlphaFold2, overlapping completely with parts of the protein covered by an experimental structure or a homology model (**Fig 3C**). Another 16.1% have no residue that overlaps with any structural information (**Fig 3C**) with the remaining pockets being partially covered with at least 1 residue found in experimental structures or homology models.

**Fig 3.**
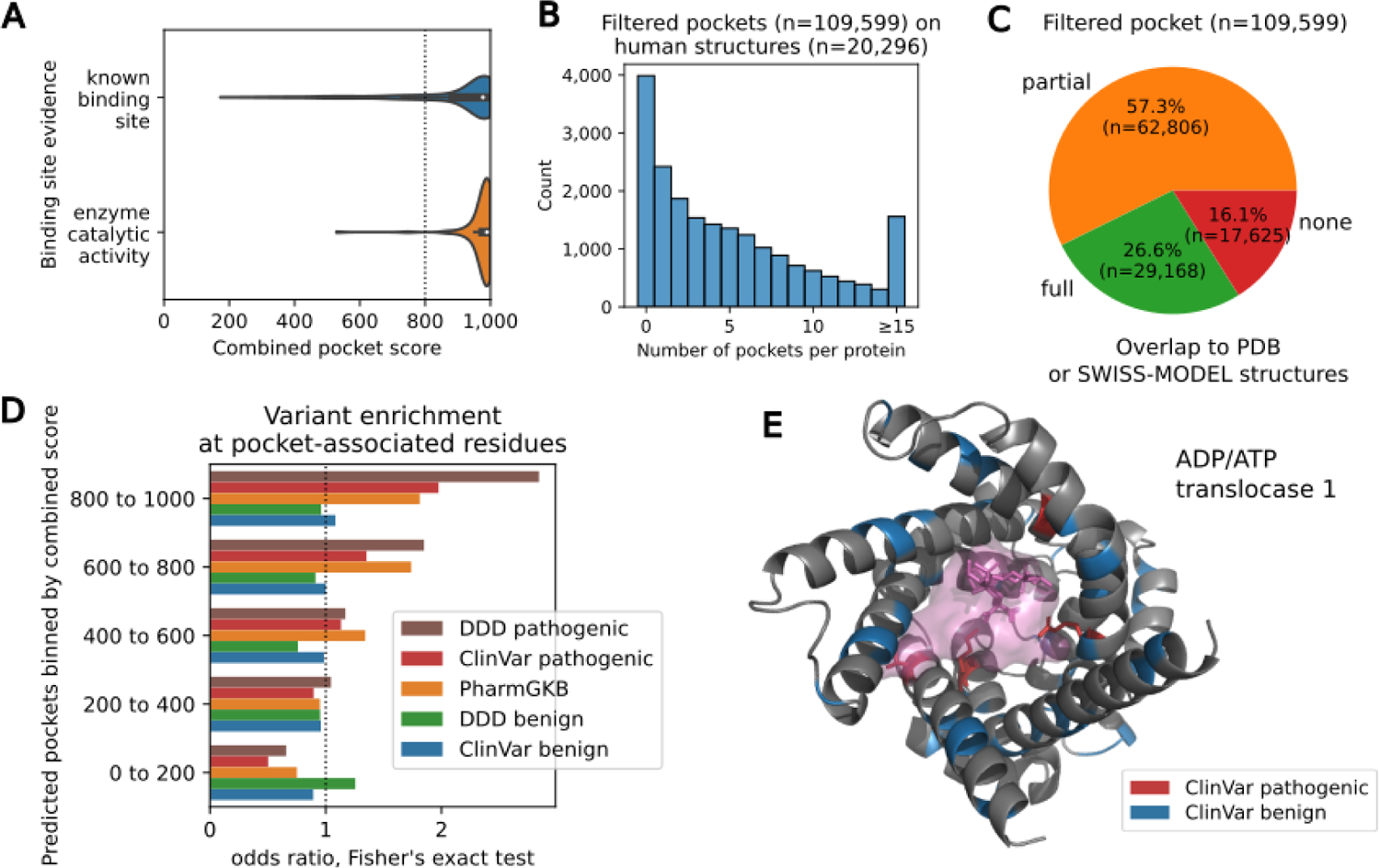
Definition of pockets, and interpretation of genomic variants. **A.** Distribution of combined pocket score at experimentally annotated binding sites, and annotated enzymes without an experimental structure. **B.** Number of filtered pockets per human structure. **C.** Coverage of filtered pocket predictions (score >800) by experimental or homology models from SWISS-MODEL. “None” corresponds to cases where no predicted pocket residue is covered by a SWISS-MODEL structure, “full” where all predicted pockets are covered by such a model and “partial” corresponding to cases where at least one residue but not all residues are covered by such a model. **D.** Enrichment of variants linked with human disorders (ClinVar, DDD) or differential response to medications (PharmGKB). Pockets were ranked into bins of pocket confidence scores and enrichments were calculated for each bin. **E.** Structure of ADP/ATP translocase 1 (P12235, ADT1_HUMAN). Residues with a pathogenic variant (red) localise at the pocket (3 of 4). Residues with a benign variant (blue) are spread across the protein (4 of 51 located in the pocket). The ligand illustrated in the protein (pink) is Carboxyatractyloside, a known inhibitor of this transporter that was added to the model with AlphaFill.

A comprehensive set of predicted pockets can be used to annotate variants associated with disease or residues linked with differences in drug responses. As expected, the protein residues in the top ranked predicted pockets show a significant enrichment in disease variants and in protein variants linked to differences in drug responses (**Fig 3D**). We find cases of disease linked genes where the pathogenic variants are specifically enriched in their pocket residues, suggesting that the driving mechanism is not simply a loss-of-function of the whole protein but of the pocket itself. For example, this is the case for the ADP/ATP translocase 1 (SLC25A4), a transporter that imports ADP into the mitochondrial matrix, exporting ATP out to the cell. In this protein we found five known pathogenic ClinVar variants, linked with mitochondrial disorders, that are found specifically at the predicted pocket site (**Fig 3E, red**) with benign ClinVar variants found away from the pocket (**Fig 3E, blue**); the ligand illustrated in the protein is Carboxyatractyloside, a known inhibitor this transporter that was added to the model with AlphaFill (Hekkelman *et al*, 2023).

### Large-scale structural modelling of human protein-protein interactions

AlphaFold2 and related methods have shown a high success rate in the prediction of structures for protein complexes (Burke *et al*, 2023; Schweke *et al*, 2023; Humphreys *et al*, 2021). To annotate residues as taking part in protein-protein interactions, we have predicted structures for 486,099 human interactions using AlphaFold2 (see Methods), generating a very large resource of predicted binding interfaces. We ranked these models according to the pDockQ confidence score (Bryant *et al*, 2022a), resulting in the identification of 108,931 predicted models of weak to high confidence (pDockQ >0.23) and 25,886 models of higher confidence (pDockQ>0.5) (**Fig 4A**). We compared the degree of experimental evidence supporting known protein-interactions with the pDockQ scores derived here. Protein interactions reported in multiple publications tend to have higher pDockQ scores (**Fig 4B**) but nevertheless, the negative set had 14.7% of models with pDockQ>0.23 and 2.3% with pDockQ>0.5 given an indication that the false-positive rates are non-negligible (**Fig 4B**). When comparing confidence scores for interactions reported in different databases we noted that protein interactions reported based on cross-linking data (xlink DB), pull-down mass spectrometry (bioplex) or a compilation of large-scale proteomics data (humap) tended to have higher fraction of high confidence models when compared with other data sources (**Fig S2A**). Overall, these results suggest that high pDockQ scores are associated with high-confidence and direct physical interactions in-line with previous observations. We next used a dataset of mutations with curated effects on protein-protein interactions (Del Toro *et al*, 2022). Using a dataset of 17,415 mutations mapped to 24,099 predicted complex structures (pDockQ>0.5) we observed that predicted interface residues are enriched for variants that cause a change in interactions relative to variants with no impact on interactions (**Fig 4C**). This enrichment was found for residues at the interface defined by a distance cut-off below 5A and it is further increased if we exclude interface residues predicted with lower confidence (pLDDT<70, **Fig 4C**).

**Fig 4.**
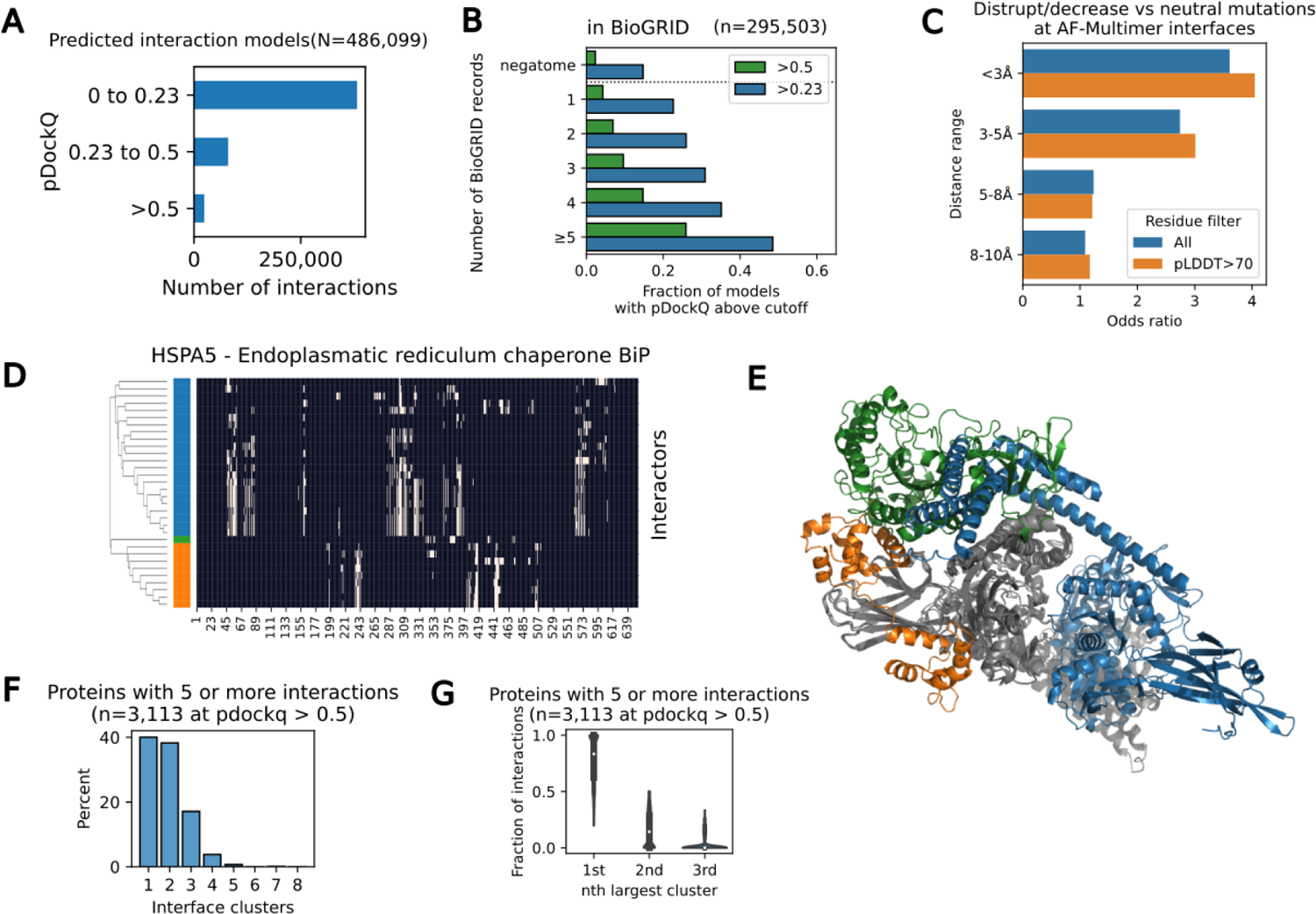
Towards comprehensive prediction of human protein interactions. **A.** AlphaFold2 predicted structures with pDockQ scores binned into poor models (pDockQ<0.23), weak models (pDockQ from 0.23 to 0.5), or median to high confidence models (pDockQ>0.5). **B.** Fraction of models with pDockQ>0.23 or pDockQ>0.5 for protein interactions supported by different numbers of independent publications as annotated in the BioGRID database. **C.** Enrichment of mutations disrupting or decreasing an interaction versus those that don’t affect the interactions at interface residues binned at different distance cut-offs between chains in the predicted models. The same enrichment is calculated and shown when including (blue) or excluding (orange) residues that are predicted to be at an interface but having low AlphaFold confidence values (pLDDT<70). D. Example of clustering of interface residues predicted between HSPA5 and 14 interactors with predicted models with pDockQ>0.5. Interface residues are annotated in the heatmap as white and the dendrogram resulting from clustering is shown to the left with the tree split into 3 selected clusters. **E.** Illustration of the AlphaFold2 predicted structure for HSPA5 in complex with 3 proteins that are representatives from the 3 different clusters found from the clustering analysis in **D.** F. Histogram of 3113 proteins, having 5 or more interactions with models with pDockQ>0.5 according to the number of interface clusters predicted based on the clustering of the interface residues. **G.** For each of 3113 proteins with 5 or more interactors selected for the analysis in F, we determined the fraction of interactors that found at the top most populated 1st, 2nd and 3rd clusters. The graph shows the distribution of fractions of proteins at these clusters with the top most populated cluster having typically the majority of the interactors.

In total, these predictions establish a putative interface for a total of 10,583 human proteins with the number of structurally resolved interactions per protein following a typical heavy-tail distribution with some proteins having a large number of predicted protein-protein interfaces (**Fig S2B**). Predicted structural models covering a large proportion of the human proteome allows for the large-scale prediction of mutually exclusive protein-protein interactions that occupy the same interface regions. To study this we selected proteins having predicted models for 5 or more interaction partners of medium to high confidence (pDockQ>0.5). For each of these proteins we clustered the predicted interface residues identified for their interaction partners identifying sets of partners occupying largely overlapping interface regions (Methods, **Fig 4D**). An example is shown for HSPA5, for which we predict models for 13 interactors (pDockQ>0.5) which were clustered into 3 different modes of interaction. Representative interactors for each of the modes are shown (**Fig 4E**) to illustrate how these can co-occur without steric hindrance. A total of 3,314 proteins had more than 5 partners modelled with a pDockQ>0.5. Clustering of the predicted interface residues suggests that most proteins have typically 1 to 3 different interaction surfaces, with a very small fraction having 5 or more different interaction modes (**Fig 4F**). Although the typical protein, with many reported interactions, tends to have multiple independent interfaces, we find that the binding partners are not predicted to be equally spread across the different interfaces but proteins tend to have one dominant interface region (**Fig 4G**).

The large collection of structural models for human protein interactions represents a resource that can be applied to the study of interface variants and other applicants. We next exemplify the application of this resource to the study of interaction-mediated control of protein abundances.

### Interaction mediated control of protein abundance levels

A significant fraction of the human proteome has been shown to be under strong post-transcriptional control whereby a change in gene expression is not propagated to protein level changes (Sousa *et al*, 2019; Gonçalves *et al*, 2017). One mechanism that has been proposed for this is that protein subunits of stable protein complexes can have a higher degradation rate when they are not bound to a complex. In this way the complex formation acts as a buffering mechanism to variation in gene expression (Gonçalves *et al*, 2017; Taggart *et al*, 2020; Dephoure *et al*, 2014). This mechanism can also explain cases where changes in protein level of a rate-limiting complex subunit can result in changes of their interaction partners, resulting in protein quantitative trait loci associations found in trans (trans-pQTL) (Gonçalves *et al*, 2017; Chick *et al*, 2016). We took advantage of the resource of structural models to investigate the structural properties of interaction mediated trans-pQTLs. We used the multi-omics cancer datasets and a gene copy-number association model to identify 727 interacting pairs, from those with structural predictions, where gene copy-number changes of a complex member predict the protein levels of its interacting partner (**Fig 5A** and **5B**, see Methods). An example of this is the TRMT6-TRMT61A interaction, with copy-number changes of TMRT6 correlating with protein level changes of both TRMT6 itself and TRMT61A (Fig 5C).

**Fig 5.**
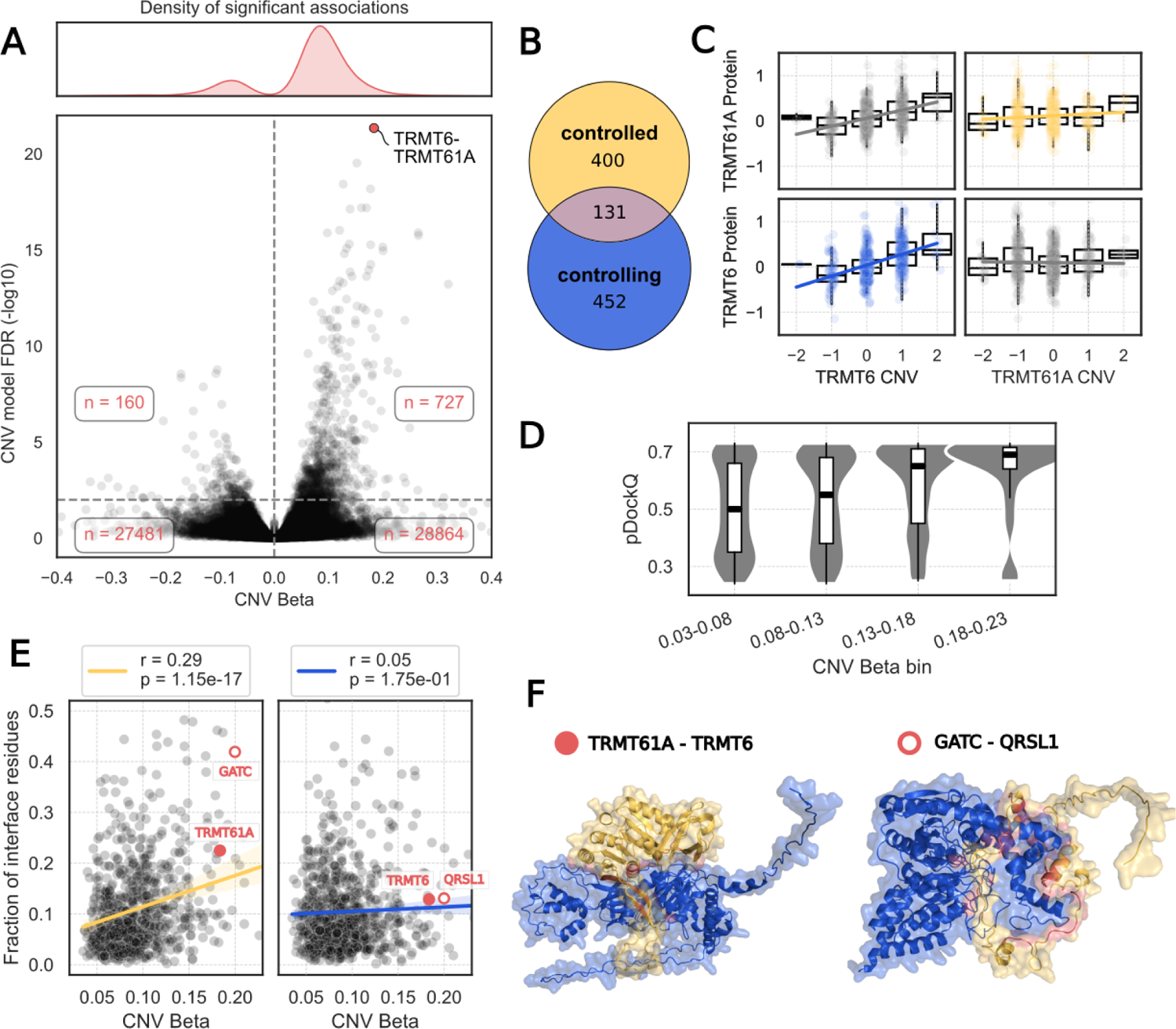
Interaction mediated control of protein abundance. **A.** Lower panel: Volcano plot of the regression coefficient of CNV_X_ (x-axis) and the associated FDR of the alternative model featuring CNV_X_ (y-axis). Each point represents an interaction, which is significant above the dashed horizontal line (FDR < 1%). Upper panel: Density of significant associations. 82% of significant associations have a positive CNV Beta. **B.** Number of unique proteins in significant associations with positive CNV Beta, by control status. **C.** Example for interaction-mediated abundance control of TRMT61A by TRMT6. TRMT6 CNV is significantly associated with both the protein abundance of itself (blue: r=0.43, p=5.7×10-31) and of its interaction partner TRMT61A (grey, top left: r=0.38, p=7.5×10-23), whereas TRMT61A CNV is neither associated with the protein abundance of TRMT6 (grey, bottom right: r=-0.02, p=0.6), or itself (yellow: r=0.08, p=0.04). **D.** Binned pDockQ distributions for significant associations with positive CNV Betas. High CNV Betas are associated with high confidence interface predictions (r=0.15, p=3.1×10-5). **E.** Strength of attenuation (CNV Beta) is positively associated with the fraction of protein residues at the interaction interface in the controlled subunit (yellow, left) but not in the controlled subunit (blue, right). **F.** Predicted structural models of two representative interactions TRMT61A-TRMT6 (pDockQ=0.73) and GATC-QRSL1 (pDockQ=0.72). Controlling proteins (TRMT6, QRSL1) are shown in blue, controlled proteins (TRMT61A, GATC) in yellow and the interaction interface residues in red. In both cases, the number of residues involved in the interaction is about equal among subunits (TRMT61A/TRMT6: 69/68 residues, GATC/QRSL1: 58/72 residues) with the controlling subunit being considerably larger (TRMT61A/TRMT6: 289/497 residues, GATC/QRSL1: 136/528 residues).

We first noted that higher confidence estimates (pDockQ score) of the AlphaFold2 model are associated with a higher degree of interaction-mediated control of protein levels for the controlled subunits (**Fig 5D**). This suggests that interactions that are more likely to be direct and/or having a larger interface are more likely to mediate these trans-pQTL effects. We then selected models of pDockQ>0.23 and observed a correlation between the degree of control and the fraction of the protein residues that is buried in the interface (r=0.29, p-value=1.15×10^-17^, **Fig 5E**) for the controlled subunit. No significant correlation was observed between the degree of control and the fraction of residues at the interface of the controlling subunits (**Fig 5E**). The result is robust to the pDockQ cut-off used (**Fig S3A**). This result is consistent with a model where there is an avoidance of leaving exposed proteins with many hydrophobic residues at the surface. Consistent with this model, we noted that interfaces of pairs that display this interaction mediated control of protein levels have a higher fraction leucine and valine (hydrophobic residues) and a lower fraction arginine (polar residue) (**Fig S3B**).

### A resource of mechanistic impacts of protein missense variants

We have generated here a comprehensive resource of annotations for protein residues and variants that may impact on protein structural stability, protein-protein interactions or interactions with small-molecules. Based on this resource we can describe the main mechanisms of pathogenicity observed in variants detected in patients. As an example application we considered variants deposited in ClinVar, of which 4.2% are currently annotated as (likely) pathogenic and 8.2% are annotated as (likely) benign (**Fig 6A**). We used AlphaMissense to predict their pathogenicity and then used our annotations to predict their most likely mechanism of action. AlphaMissense predicted 24.7% as pathogenic and 65.4% as benign, with most known annotations from ClinVar correctly predicted. From all of the putative pathogenic predictions our resource infers that the primary mechanism of effect was stability changes (39.5%), followed by pockets (14.8%) and protein-protein interfaces (9.2%), with the remaining unclassified. ClinVar primarily contains variants that are found in patients of rare disorders and therefore it is expected that the main impact of such variants would be loss-of-function through destabilisation. We compared these proportions with other lists of variants linked with human phenotypes, including variants found in the Deciphering Developmental Disorders (DDD) project, recurrent mutations in cancer (Cancer Hotspots), variants with a known impact on interactions curated by IntAct, and variants linked with differences in response to medications (PharmGKB). We selected from each list of variants those that AlphaMissense predicts as pathogenic and then we assigned a putative mechanism using our resource (**Fig 6B**). The cancer associated variants and those known to impact in interactions had the largest fraction of variants at putative protein-protein interfaces and PharmGKB had the largest fraction of variants at putative pocket residues (**Fig 6B**). Nevertheless, many variants had impacts that are likely mediated by protein stability effects (**Fig 6C**).

**Fig. 6.**
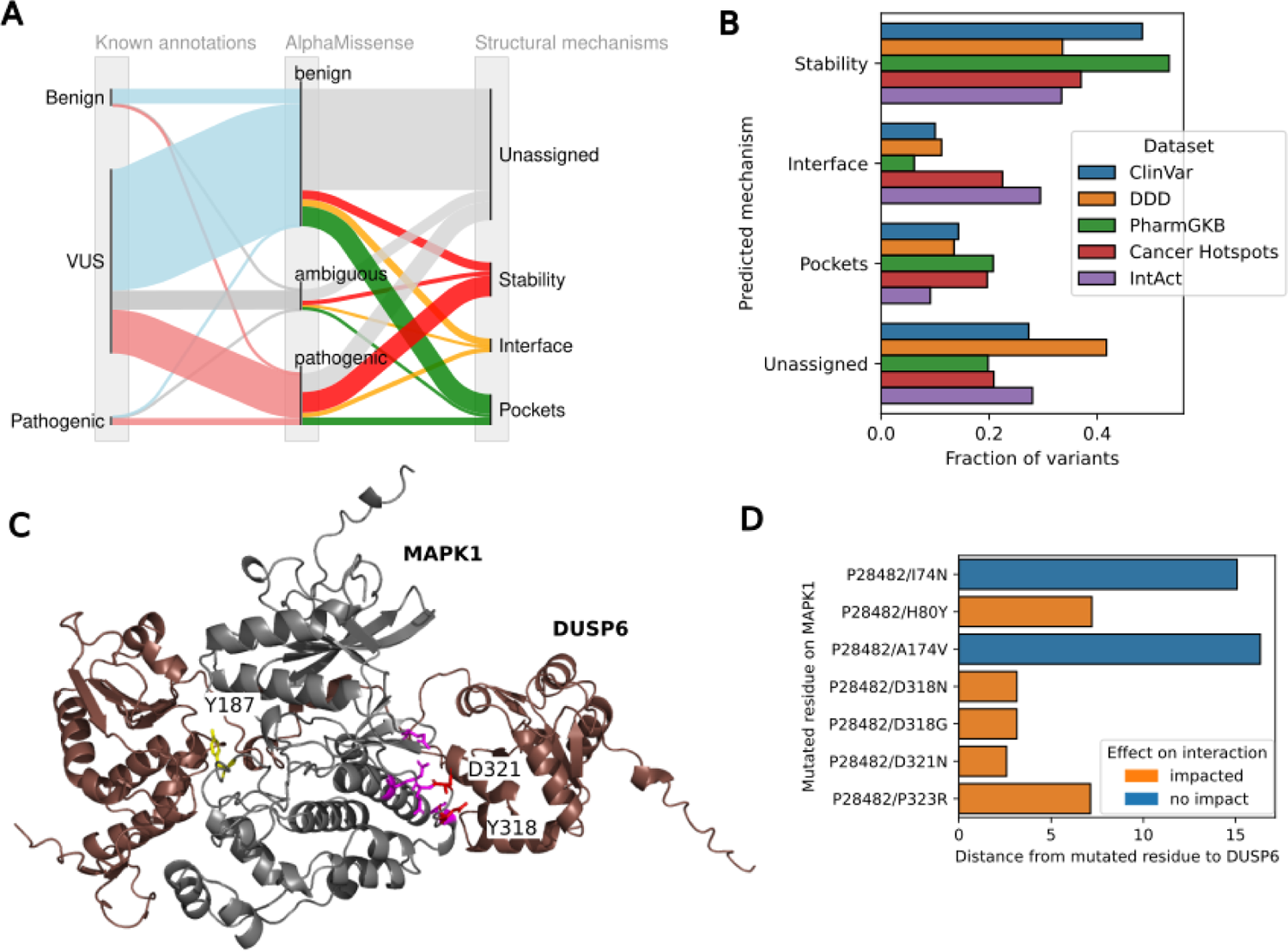
Putative mechanisms in different (pharmaco)genomic data sets. **A.** Sankey plot annotating ClinVar missense variants with pathogenicity scores from AlphaMissense and a likely structural mechanism using our resource **B.** Distribution of putative mechanisms of variant effects across different sets of disease linked or drug-response linked variant datasets **C.** MAPK1-DUSP6 AF-Multimer predicted structure highlighting relevant residues in MAPK such as the Y187 phosphorylation site (yellow) that is a known target site for the phosphatase, 2 interface mutations annotated to disrupt binding (D231 and Y318 in red) and 4 residues with known pathogenic clinical variants that are predicted to be at the interface (pink residues). **D.** Distances between mutated residue on MAPK1 and DUSP6 with annotated impact on binding

The combination of predicted structural models and known impacts of mutations from different data sources can help validate the predicted structures and shed light on the disease mechanisms. As an example we have identified 12 interactions with: a predicted structural model of high confidence (pDockQ>0.5); at least 2 mutations in predicted interface residues known to disrupt binding; and at least 2 mutations at predicted interface residues known to cause a clinical disorder. An example of this is the interaction between MAPK1 and DUSP6 with a predicted structure shown in **Fig 6C** (pDockQ=0.62). While there is no experimental structure for this complex, there are experimental structures of homologous domains. This predicted model places the N-terminal region of DUSP6 at a docking site in MAPK1, in a binding mode that is, in part, similar to the complex formed between p38α and the MAPK phosphatase 5 (MKP5) (Zhang *et al*, 2011). The catalytic domain of DUSP6 is oriented such that it can act on the known target phosphosite at tyrosine 187 of MAPK1 (**Fig 6C**, yellow). The 2 mutations curated in IntAct, for MAPK1 that result in decrease of binding to DUSP6 are predicted to be at the interface (**Fig 6C**, Y318 and D321). These combined observations strengthen the validity of the predicted model which can then be used to assign a mechanism to other clinical variants. In this case, we have identified 4 MAPK1 disease-causing variants at the interface regions (**Fig 6C**, pink) indicating that loss of this interaction is a relevant disease mechanism. In support of this, a report has identified several mutations in patients with a neurodevelopmental disease that are likely to act via the disruption of this interface, resulting in increased activation of the kinase (Motta *et al*, 2020). This model places the mutations with the strongest impact on the interaction at or near the interface (**Fig 6E**).

We illustrate how structure-based prediction of variant effects offers some insight into the mechanisms of disruption of protein function. The residue level annotations and proteome-wide prediction of variant effects are provided for public use at the ProtVar resource (www.ebi.ac.uk/ProtVar/) (Stephenson *et al*, 2024) where any human protein variant can be queried.

## Discussion

There are millions of missense variants found in the human population that lack annotations. Here, we have taken advantage of AlphaFold2 predicted structures to annotate variants according to their impact on protein stability, protein interfaces and small-molecule binding sites.

In our analysis of cancer samples we find an expected link between predicted destabilising variants and decrease of protein abundances. However, the effect size of this association is very modest and only clear in cases where the gene is in a single copy. Our analysis provides examples where predicted loss of stability coincides with different degrees of changes in protein levels. This lack of signal could be due to many unaccounted factors, including the interaction with chaperones and other protein quality control mechanisms. In addition, the samples studied are highly variable in their genetic background and other factors. Experimental studies of such variants in isogenic backgrounds would be required to validate their consequences and to study the cellular mechanisms that relate stability with degradation.

We generated a large resource of predicted structures for human protein complexes and illustrate the use of this resource to identify which interaction partners may be mutually exclusive as they would occupy the same interface. It has been noted that such predictions may have a higher rate for paralogs (Burke *et al*, 2023). This may be due to the fact that paralogs can often be included in the same MSA, and AF2 will not easily distinguish the evolutionary signals that drive a prediction to a specific interface. Despite potential errors, we identified interface regions for 10,583 proteins (pDockQ>0.5) and we suggest that interaction regions within these proteins may often be correct even if many structures for complexes are incorrect. In other words, even low confidence structures may be useful to identify regions within a monomer that may be interacting residues. We note also that the negative control interactions have a non-negligible error rate (2.3% with pDockQ>0.5) and protein interactions supported by multiple experimental evidence tend to have a larger fraction of models of higher confidence. Given the sparse nature of protein-protein interactions inside cells, these error rates are expected to result in large numbers of false-positives when modelling random pairs or protein-pairs with weak experimental evidence for interactions.

Nearly 80% of the predicted protein complex structures resulted in a model of very low confidence indicating that only a fraction of the total computational time went into predicting models of high confidence. Proteome-wide structural models of protein complexes will require efforts into efficient predictions of pairs of proteins that are likely to result in high confidence structural models. Such efficient models could be used to prioritise pairs for more computationally demanding structural modelling. In addition, we have focused here on modelling pairs of proteins and additional work will be required to predict structures for larger assemblies (Bryant *et al*, 2022b). The models provided here should be useful for training of machine learning models for diverse tasks, including models that can prioritise protein-pairs for future protein complex predictions and for training of AlphaFold-like models.

Destabilisation was the major mechanism assigned to ClinVar variants that are predicted to be deleterious by AlphaMissense. This is not unexpected since loss-of-stability is also the most probable loss-of-function missense variant produced from a random set of variants. However, despite the fact that we obtain a very comprehensive set of interface models, we only cover around half of the human proteome and only a small fraction of the universe of human-human protein interactions. So a higher fraction of interface disrupting variants would be mapped with a more comprehensive set of interaction models. Nevertheless, we could relate datasets of variants from recurrent mutations and those linked with differences to drug response with differences in proportions of annotated mechanisms. Recurrent cancer mutations were more likely to be linked with protein interfaces and those associated with differential drug responses had a higher fraction annotated to small-molecule binding pockets.

In summary, we established here a large resource of annotations for the impact of protein missense variants. We illustrate several applications of this resource and make all of the models, annotations and pre-computed variant effects accessible via the ProtVar resource (www.ebi.ac.uk/ProtVar/).

## Methods

### Stability and Pocket predictions

Alphafold2 structures were pre-processed prior to stability predictions by removing low-confidence regions. These regions were defined as having an average pLDDT<50 over a 10 amino acid window. FoldX stability predictions were then performed using the BuildModel command for all possible mutations at all positions in the structure. The AFDB model for FGFR1 was additionally processed to remove incorrect domain interactions due to the lack of TM in the model. We rebuilt the FGFR1 model based on the domain-domain orientations of FGFR2 X-ray structure (1E0O) which was also trimmed to remove low-confidence regions and used for stability predictions using FoldX.

We used AutoSite with default parameters to obtain raw pocket predictions for AlphaFold2 structures. We benchmarked different pocket metrics for their ability to predict known binding sites from BindingMOAD. We found that a version of the AutoSite empirical composite score modified to include mean pLDDT of pocket-associated residues performed best (Fig S2). For testing generalisability, we adapted the approach from (Akdel *et al*, 2022) where we considered as true positives pockets found in established enzymes that had no known experimental structures or homology models available in SWISS-MODEL.

Pharmacogenomic variants were downloaded from pharmgkb (Whirl-Carrillo *et al*, 2012, 2021) To test for enrichment at binding sites, we defined pocket-associated residues (<4.5A). For global enrichment, We then tested whether positions with a pathogenic variant were enriched in all pocket-associated residues (within the specific score of interest) compared to all other residues in human single-fragment AF2 structures. For enrichment within a particular pocket, we similarly tested all positions with a variant against enrichment against the rest of the structure. Resulting p-values were FDR-corrected.

Variants from ClinVar (2024-04-16) were mapped to the proteome using ProtVar, and filtered to keep missense variants with at least one review star. Sankey plots were obtained using floWeaver (Lupton & Allwood, 2017).

### Prediction of structures for protein complexes and interface analyses

We used Alphafold2-Multimer (Evans *et al*, 2021) to predict the structures of binary protein-protein interactions which have not been previously built using the FoldDock pipeline (Burke *et al*, 2023). The standard sequence databases and tools as defined in the AF2 pipeline were used to search for homologous sequences. Multiple sequence alignments were generated using default values for all parameters. Initial models were built using the max_recycles=3 and maxpdb=20 options of AF2. For structures with a low pDockQ score, subsequent models were built using a higher number of max_recycles (10, 20 and 50). Structures were post-processed using the FoldX (Delgado *et al*, 2019) RepairPDB command. Where multiple models were built for a pair of proteins, the model selected for analysis was that which has the highest of pDockQ.

### Multi-Omics Data Collection and Processing

For our analysis, we used multi-omics cancer data made available by the CPTAC consortium that was compiled and standardised as described in (Sousa *et al*, 2023). We excluded samples from the children’s brain tumour tissue consortium (cbttc) and colorectal tissue (tcga-coread) batches due to low coverage of proteomic measurements. The remaining dataset contained 651 patient-derived cancer samples with matched somatic mutations, copy number variation (CNV), mRNA and protein expression data from six different tissues: liver (hcc-proteogenomics), ovarian (tcga-ov), kidney (discovery-ccrcc), uterus (discovery-ucec), colon (colon-opportunities), and breast (tcga-brca). The CNV data was obtained as discretized GISTIC2 scores. A discretized score of ™2 represents a strong copy number loss (homozygous deletion), ™1 a shallow deletion (heterozygous deletion), 0 the diploid state, 1 a low-level gain of copy number (general, broad amplification) and 2 a high-level increase in copy number (focal amplification). Protein and mRNA expression data were normalised using a linear regression to remove the confounding factor related to the experimental batch and converted to log fold changes by dividing the batch-corrected abundance by the median of the experimental batch. Due to the sparseness of the protein data, we selected 8,791 genes that had at least 25% sample coverage in each of the three omics modalities (CNV, mRNA, protein).

### Variant Effect Predictions and Protein Abundance

Some genes featured more than one somatic mutation in a sample. In such cases, we only retained the mutation with the highest AlphaMissense score. We identified and removed six hypermutated samples carrying more than 500 mutations. We then removed 1,465 genes from the analysis which had little concordance of mRNA and protein levels (Pearson’s r < 0.3), assuming that in these cases, there were either technical or biological factors preventing genetic changes to be detectable on the protein level. Next, we wanted to calculate the residual variance in protein abundance after accounting for mRNA abundance. We first fitted a linear model with the protein abundance *P*_*Y*_ as the dependent variable, mRNA abundance *T*_*Y*_ as the independent variable, the experimental batch as a covariate *C* and error term ε (equation 1). Here, *C* is the matrix containing the dummy variables for the different experimental batches and β_c_ is the vector of coefficients for the dummy variables in *C*. After fitting the linear model, we derived the protein residuals *P*_*Y*′_ by calculating the predicted protein abundance *P*_*Y*_ using the estimated parameters β_0_, β_1_ and β_c_ and subtracting that from the measured protein abundance *P*_*Y*_ (equation 2).

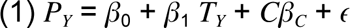

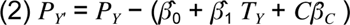

The residual variance *P*_*Y*′_ could then be compared between different variant effect predictions. For this, we had to discretize the VEP scores: AlphaMissense scores smaller than 0.34 were classified as benign, larger than 0.564 as pathogenic, and in between as ambiguous. FoldX scores smaller than ™2 were classified as stabilising, larger than 2 as destabilising, and in between as neutral.

### Nested Linear Models for the Identification of Protein Associations

To identify interactions where protein *P*_x_ can influence the abundance of protein *P*_*Y*_, we asked whether including the abundance of *P*_x_ next to the transcript level of *P*_*Y*_(*T*_*Y*_) would help in explaining the detected protein abundance of *P*_*Y*_. To only capture directed interactions, we opted to use *CNV*_*x*_ as a proxy for the protein abundance *P*_x_ instead of *P*_x_ directly, because there is no cellular mechanism by which *P*_*Y*_could influence the level of *CNV*_*x*_. In more technical terms, we employed two nested linear models for each pair of interacting proteins. The null model M 0 only included *T*_*Y*_as an independent variable (with experimental batch C as covariate), and *P*_*Y*_as a dependent variable (equation 3). The alternative model included *CNV*_*x*_as an additional independent variable (equation 4).

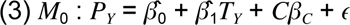

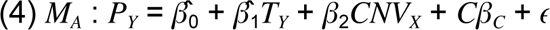

Using a likelihood ratio test (LRT), we then asked whether the alternative model would improve the goodness of fit in comparison to the null model (equation 5). The LRT statistic follows a chi-squared distribution under the null model, allowing us to derive a p-value for the null hypothesis using equation 6. We then adjusted the p-values for multiple testing using the Benjamini-Hochberg procedure, resulting in the false discovery rate (FDR). For the subsequent analyses, we used an FDR < 0.01 as a threshold to identify protein-protein interactions where CNV X helps explain P Y.

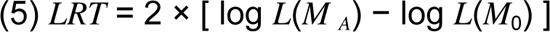

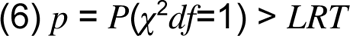

### Characterization of FGFR1 patient variants

#### Cell Culture

GT1-7 mouse hypothalamic GnRH neuronal cells were obtained from the Sigma-Aldrich and cultured in Dulbecco’s Modified Eagle’s Medium (DMEM) supplemented with 10% fetal bovine serum (FBS), 100 U/ml penicillin, and 100 µg/ml streptomycin (Thermo Fischer Scientific, Waltham, MA, USA). The cells were maintained at 37°C in a humidified atmosphere containing 5% CO2. The culture medium was changed every two days, and cells were passaged upon reaching 70-80% confluency.

Plasmid Constructs: HA-tagged plasmids encoding wild-type (WT) and mutant variants of FGFR1 were acquired from Genscript (Singapore). Prior to their use in transfection experiments, the integrity and sequence of these constructs were confirmed through DNA sequencing to ensure accuracy and reliability.

#### Transient Transfection and FGF2 Treatment

GT1-7 cells were transiently transfected with plasmids encoding either wild-type (WT, transcript ENST00000425967) or mutant FGFR1 using Lipofectamine 3000 reagent (Thermo Fischer Scientific, Waltham, MA, USA), following the instructions provided by the manufacturer. Initially, cells were plated in 6-well plates at a density of 2.5 x 10^5 cells per well and allowed to adhere for 24 hours. For transfection, 2 µg of plasmid DNA was combined with 5 µl of Lipofectamine 3000 reagent in Opti-MEM Reduced Serum Medium (Invitrogen) and incubated for 15 minutes at room temperature to form DNA-lipid complexes. These complexes were then added to the cells, and the plates were gently swirled to ensure uniform distribution. After 6 hours, the medium containing the transfection reagent was replaced with fresh complete DMEM. The cells were then cultured for an additional 36-48 hours before subsequent experiments. For FGF2 treatment, cells were stimulated with 100 ng/ml FGF2 for 60 minutes after the transfection period.

#### Cell Growth Assay

GT1-7 cells were transiently transfected with plasmids encoding wild-type (WT) and mutant FGFR1. Cell growth was assessed using the MTT assay kit (Abcam, USA) according to the manufacturer’s instructions. At specific time points post-transfection, MTT reagent was added to the cells, and after incubation, the absorbance was measured at 570 nm using a microplate reader. The absorbance values, indicative of viable cell numbers, were used to evaluate the impact of FGFR1 variants on GT1-7 cell proliferation.

#### Protein Extraction and Cell Surface Protein Isolation

After 36-48 hours of transfection, cells were lysed using RIPA buffer to extract total proteins. For the isolation of cell surface proteins, we utilized the Pierce Cell Surface Protein Isolation Kit (Thermo Fischer Scientific, Waltham, MA, USA) following the manufacturer’s instructions. The cells were initially cooled at 4 °C for 15-20 minutes, followed by biotinylation using EZ-Link Sulfo-NHS-SS-Biotin for 30 minutes at 4 °C. The biotinylation reaction was halted with a quenching buffer, and cells were subsequently lysed on ice using the kit’s lysis buffer. Biotinylated proteins were then isolated by incubating the lysates with NeutrAvidin Agarose. After three washes, the proteins were eluted using the kit’s elution buffer, which was heated to 95°C for 5 minutes.

#### Western Blotting

The extracted total and cell surface proteins were separated on a 4–12% Nupage gel and then transferred to a Polyvinylidene Fluoride (PVDF) membrane (Thermo Fischer Scientific, Waltham, MA, USA). The membrane was blocked using a non-fat milk buffer and incubated with the appropriate primary antibody overnight at 4 °C. Subsequently, the membrane was incubated with an HRP-conjugated secondary antibody. Protein bands were visualized using the Chemdoc system (Bio-Rad Laboratories, Hercules, CA, USA) and quantified with ImageJ software (NIH, Bethesda, MD, USA).

### Data and code availability

All of the predictions for stability, pockets and AlphaFold2 multimer models can be accessed via the web interface at https://www.ebi.ac.uk/ProtVar/ and are available for download in bulk at https://ftp.ebi.ac.uk/pub/databases/ProtVar/. Code used for this project is available at https://github.com/jurgjn/af2genomics/

## Supplementary Figures

**Fig S1.**
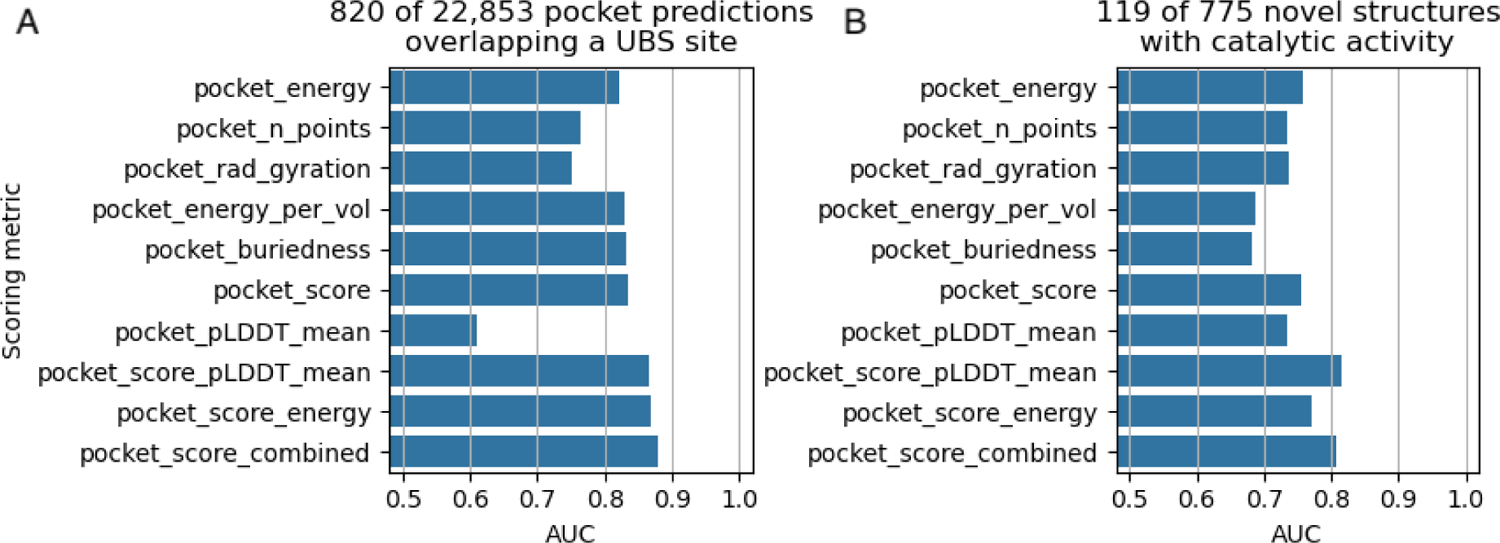
Recall of known binding sites (top) and enzymatic activity (bottom) using different pocket prediction scoring metrics. pocket_energy, pocket_n_points, pocket_rad_gyration, pocket_energy_per_vol, pocket_buriedness and (AutoSite) pocket_score are defined as in (Ravindranath & Sanner, 2016); pocket_pLDDT_mean is the mean pLDDT values of pocket-associated residues (within 4.5A of the pocket). pocket_score_pLDDT_mean, pocket_score_energy and pocket_score_combined are modifications of the AutoSite score defined as follows: score = n_points * buriedness**2 / radius_gyration score_pLDDT_mean = n_points * buriedness**2 * pLDDT_mean**2 / radius_gyration score_energy = energy * buriedness**2 / radius_gyration score_combined = energy * buriedness**2 * pLDDT_mean**2 / radius_gyration

**Fig S2.**
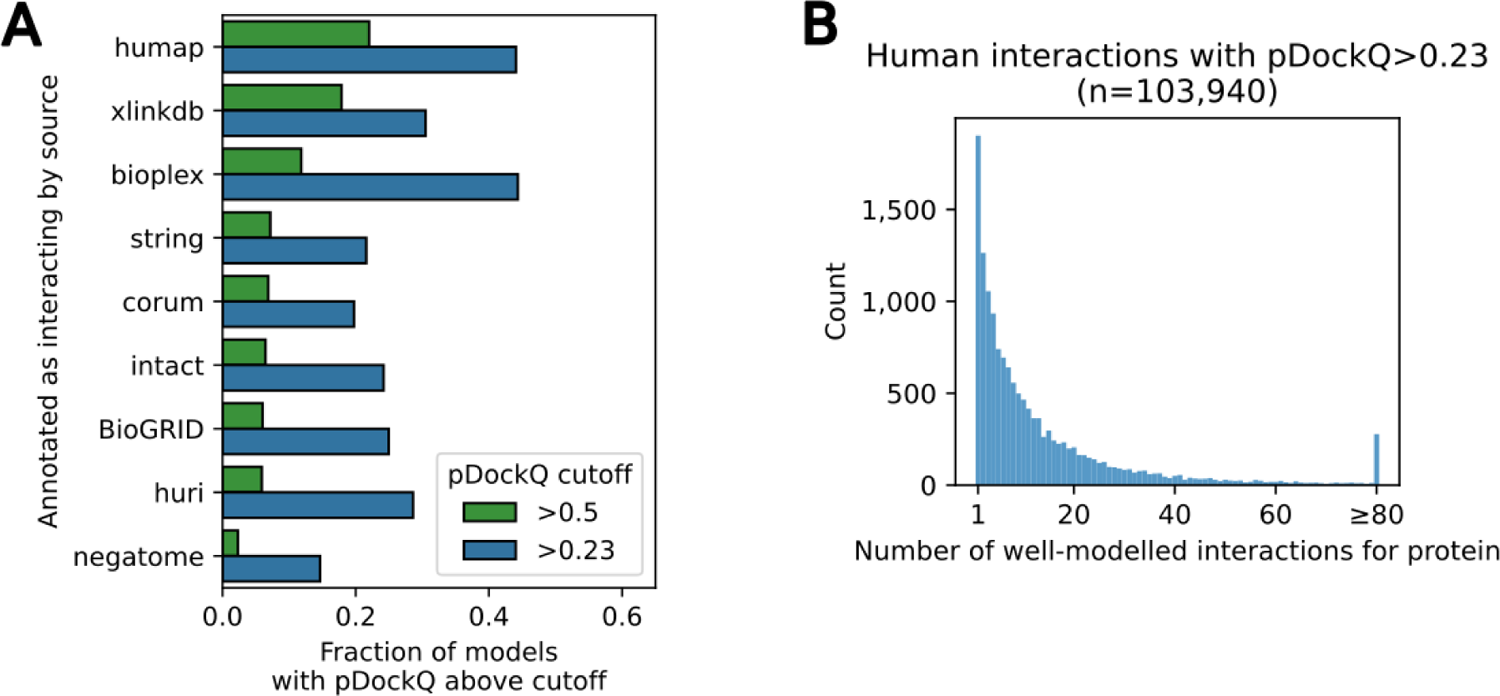
Predicted models per interaction type and binned per protein. **A.** pDockQ confidence scores for interactions reported in different databases. A higher fraction of confident models (pDockQ>0.5) was observed for interactions derived by affinity-purification (humap, bioplex) and cross-linking data (xlinkdb) than for yeast-two-hybrid (huri) or general compilations of data (BioGRID, intAct). **B.** Number of models with pDockQ>0.23 per protein. The distribution of the number of models per protein follows a heavy tail distribution that is similar to the distribution of the number of experimentally determined protein-protein interactions per protein. The majority of proteins having few (<15) interaction models with a long tail of proteins having a large number of models.

**Fig S3.**
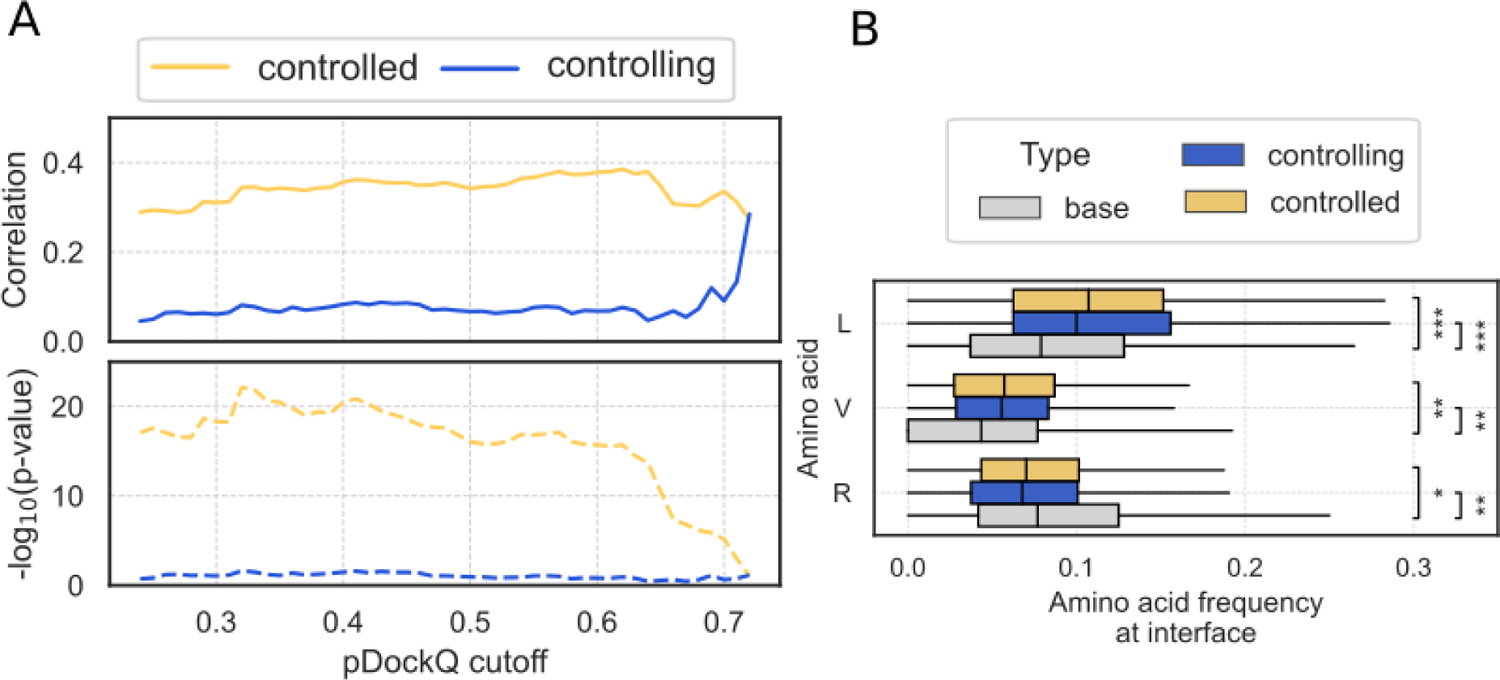
Relation protein surface properties and interaction mediated control of protein abundance. **A.** Correlation (upper plot) and significance of correlation (lower plot) between the strength of attenuation (CNV Beta) and the fraction of protein residues at the interaction interface in the controlled subunit (yellow) and in the controlling subunit (blue) for different pDockQ cutoffs. Enrichment of leucine and valine residues and depletion of arginine residues at the interfaces where there is a strong predicted control of protein levels by protein-protein interactions.

## Acknowledgements

This research was funded by a grant from the National Institutes of Health to N.J.K (Psychiatric Cell Map Initiative (PCMI); U01MH115747) and the Quantitative Biosciences Institute (QBI) at UCSF. P.B. is supported by the Helmut Horten Stiftung and the ETH Zurich Foundation. This work is supported by European Molecular Biology Laboratory core funds and Open Targets.

